# SCCNV: a software tool for identifying copy number variation from single-cell whole-genome sequencing

**DOI:** 10.1101/535807

**Authors:** Xiao Dong, Lei Zhang, Xiaoxiao Hao, Tao Wang, Jan Vijg

**Author notes:** Emails*: XD;, LZ, XH, TW, JV.

## Abstract

**Background:** Identification of de novo mutations from cell populations requires single-cell whole-genome sequencing (SCWGS). Although many experimental protocols of SCWGS have been developed, few computational tools are available for downstream analysis of different types of somatic mutations, including copy number variation (CNV).

**Results:** We developed SCCNV, a software tool for detecting CNVs from whole genome-amplified single cells. SCCNV is a read-depth based approach with adjustment for the whole-genome amplification bias.

**Conclusions:** We demonstrate its performance by analyzing data collected from most of the single-cell amplification methods, including DOP-PCR, MDA, MALBAC and LIANTI. SCCNV is freely available at https://github.com/biosinodx/SCCNV.

## Background

Each single cell in a tissue or cell population has its own unique genome due to accumulating de novo mutations, such as single-nucleotide variations (SNVs), structural variations (SVs), copy number variations (CNVs) and aneuploidies. The frequency and spectrum of the mutations reflect the loss of genome integrity of a cell population, important to cancer and aging [1]. To detect the mutations unique to a single cell, single-cell whole-genome sequencing (SCWGS) is necessary. However, SCWGS requires whole-genome amplification (WGA), which often causes allele-specific or locus-specific bias. The bias essentially constrains the usage of variant callers designed for non-amplified bulk DNA. We recently developed a new software tool, SCcaller, that corrects for allelic bias using SNP markers [2]. Another major type of mutation, i.e., copy number variation (CNV), is detected on the basis of read depth, which is also affected by locus-specific amplification bias [3–5]. Yet, the only available tool is a webserver which requires uploading of the SCWGS data [6]. The uploading could be time consuming because SCWGS data is often in big size, i.e., 100 GB per single cell. This lack of a software tool for detecting CNVs is becoming a limiting factor for the application of SCWGS data for which many experimental protocols are available, e.g., DOP-PCR, MDA, MALBAC and LIANTI [2–5, 7]. To meet this need, we present SCCNV, a software tool to identify CNVs from SCWGS. SCCNV is also based on a read-depth approach; it controls not only bias during sequencing and alignment, e.g., bias associated with mapability and GC content, but also the locus-specific amplification bias. We demonstrate the application of SCCNV to SCWGS data of multiple experimental protocols, i.e., DOP-PCR, MDA, SCMDA, MALBAC and LIANTI.

## Implementation

SCCNV was written in Python. It uses SCWGS data after alignment as input (i.e., a bam file). First, it divides the genome into bins of equal size (500kb as default), and counts the numbers of reads per bins of a cell. SCCNV then normalizes mapability, which indicates the efficiency of the alignment to a genomic region. For a bin *b* of a cell, SCCNV adjusts the raw number of reads, denoted by *NR_raw_*, by dividing over the mapability *M*,

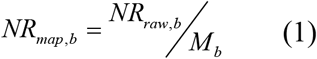

where mapability *M* is a value ranging from 0 to 1. SCCNV uses Encode Align100mer mapability score, downloaded from the USCS genome browser, and calculates the mapability of each bin by using their weighted average.

Then, SCCNV normalizes for GC content. For a cell, SCCNV calculates the percentile of GC content of each bin. For a bin *b* of the cell, its number of aligned reads after normalizing GC content, *NR_GC,b_*, is,

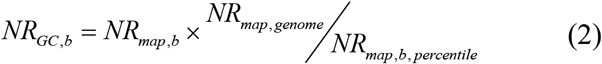

where *NR_map,genome_* is the average *NR_map_* per bin of all bins from the cell; *NR_map.b,percentile_* is the average *NR_map_* per bin of bins in the same GC percentile as bin *b*.

A consistent pattern of sequencing depth presents in different cells amplified using the same experimental protocol. This reflects the pattern of locus-specific amplification bias. Therefore, the bias is normalized across all cells in a particular batch and experiment. First, to make the *NR_GC,b_* comparable across cells, SCCNV converts it to a raw copy number estimate, denoted by *CN_raw,b_* for bin *b* of cell *c*, as follows,

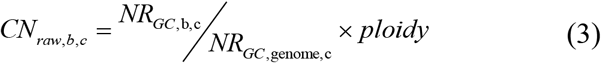

where *NR_GC,genome,c_* is the median *NR_GC,c_* per bin in the genome of cell *c*; ploidy is 2 by default. Second, the adjusted copy number is estimated as,

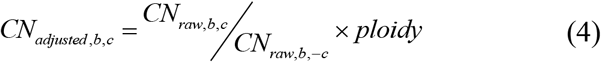

where CN_raw,b,-c_ denotes the average *CN_raw_* for bin *b* across all cells except cell *c*. Then SCCNV uses a sliding window approach to further minimize amplification noise. By default, a window includes 10 500kb bins, i.e., 5 Mb in total, with a 500kb step size between two neighboring windows.

SCCNV then models the distribution of *CN_smoothed,b,c_* of all bins in autosomes of a cell *c* as a normal distribution N(μ, σ_c_^2^). The μ=2, and σ is estimated as,

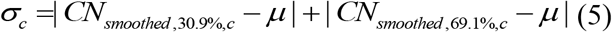

where *CN_smoothed,30.9%,c_* and *CN_smoothed,69.1%,c_* are the 30.9% and 69.1% percentiles of the *CN_smooth,b,c_* of all bins in the autosomes, respectively.

Assuming equal priors, for a bin *b* and a given possible copy number *k* ∈ {0, 1, 2, 3, 4}, its posterior probability is,

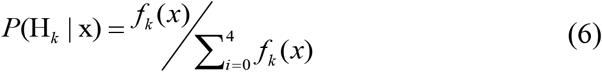

where *x* is the *CN_smoothed,b,c_*; *f_i_*(*x*) is the probability density function of a normal distribution,

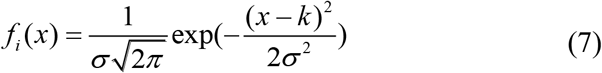

where the variance *σ*_*c*_^2^ is calculated according to equation (5).

SCCNV allows <1 false positive per cell. Therefore, it determines bin *b* as a copy number variant when,

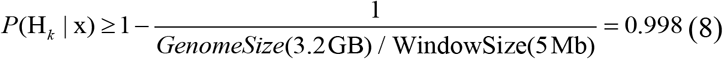

## Results

Figure 1 presents the major steps in SCCNV, including normalizing mapability and GC content, normalizing locus-specific amplification bias and inferring copy number. Before and after each step, SCCNV also generates intermediate results for users to monitor its performance. Using a single cell amplified with MDA as an example (SRA id: SRR2141574), we first show that our procedure normalized the bias in number of reads due to mapability as well as GC content (Figure 2 and 3). Second, we show that locus specific amplification bias was adjusted in Figure 4. Finally, copy number variations can be determined based on a statistic simulation and testing (Figure 5 and Equations 5-8).

**Figure 1.**
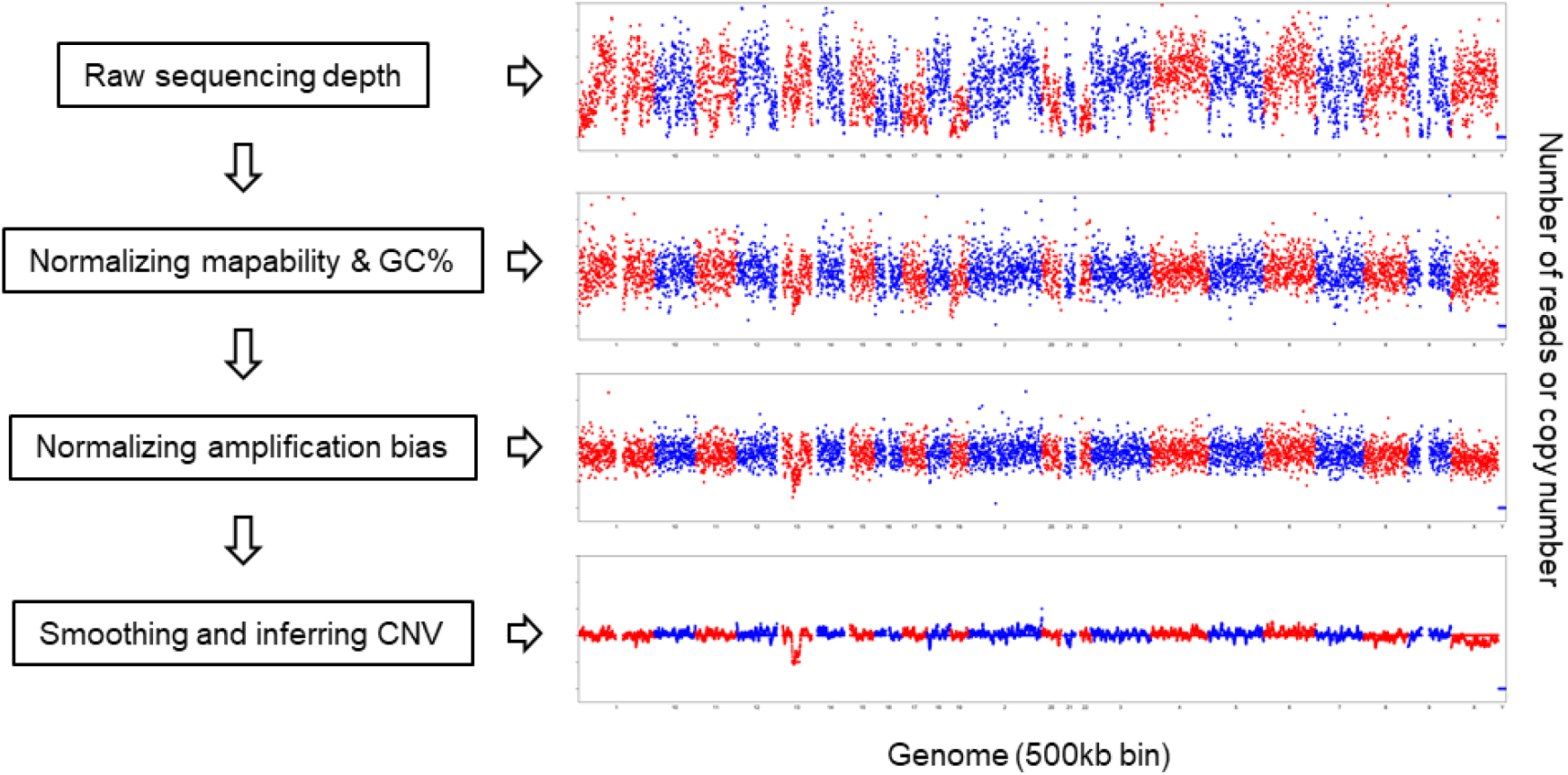
Major steps in SCCNV. The example is generated using a cell amplified with MDA (SRA id: SRR2141574).

**Figure 2.**
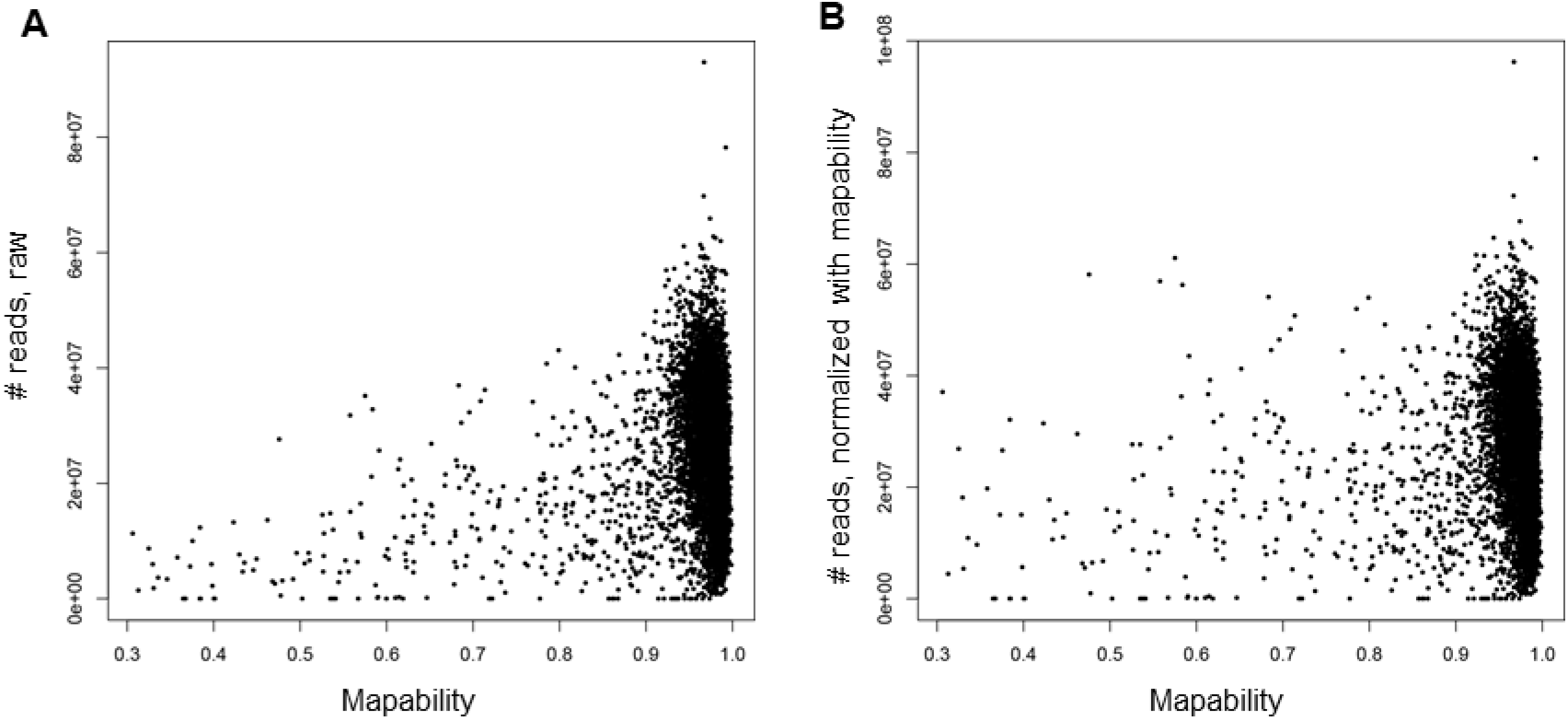
Mapability and number of reads per 500kb bin before (A) and after normalizing (B) the mapability. This example was generated using a cell amplified with MDA (SRA id: SRR2141574).

**Figure 3.**
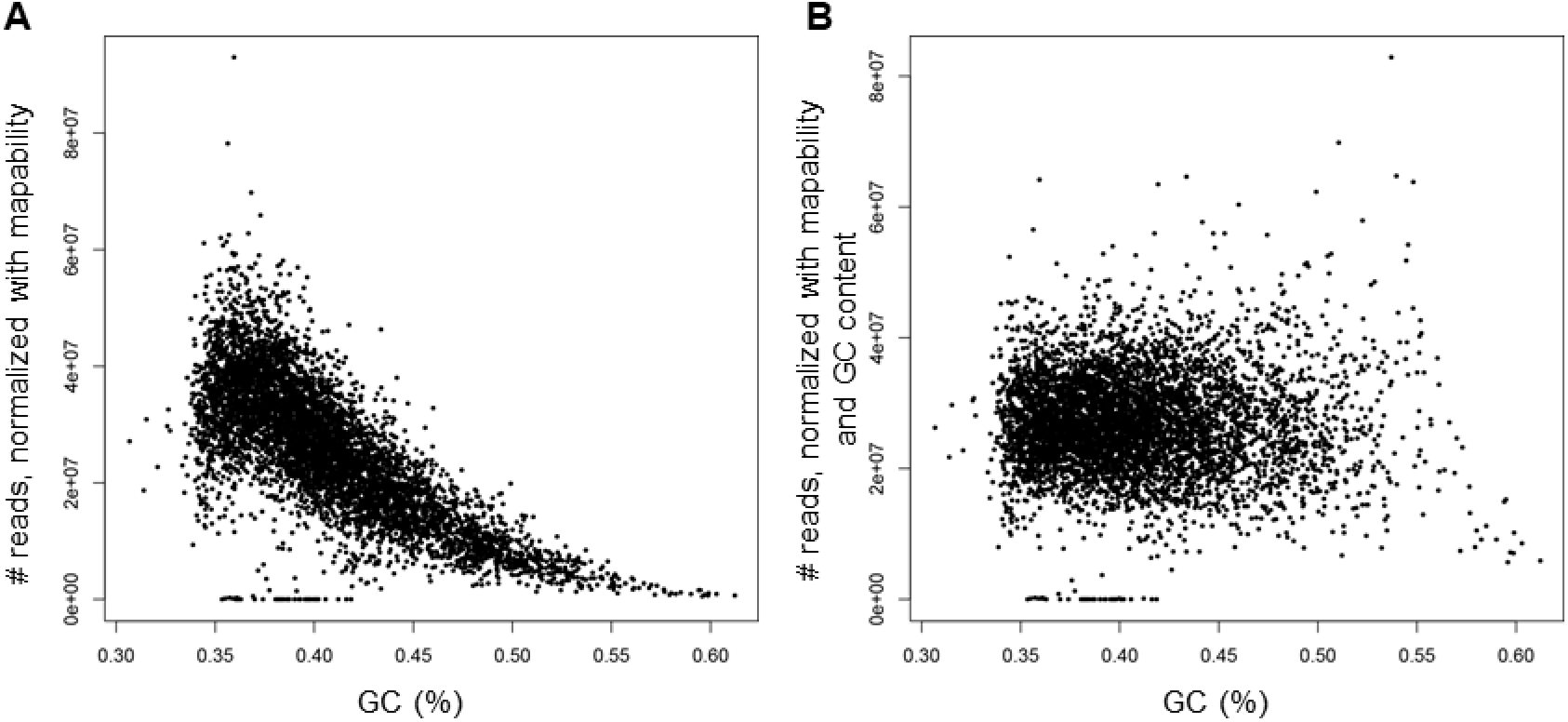
GC content and number of reads per 500kb bin before (A) and after (B) controlling for GC content. This example was generated using a cell amplified with MDA (SRA id: SRR2141574).

**Figure 4.**
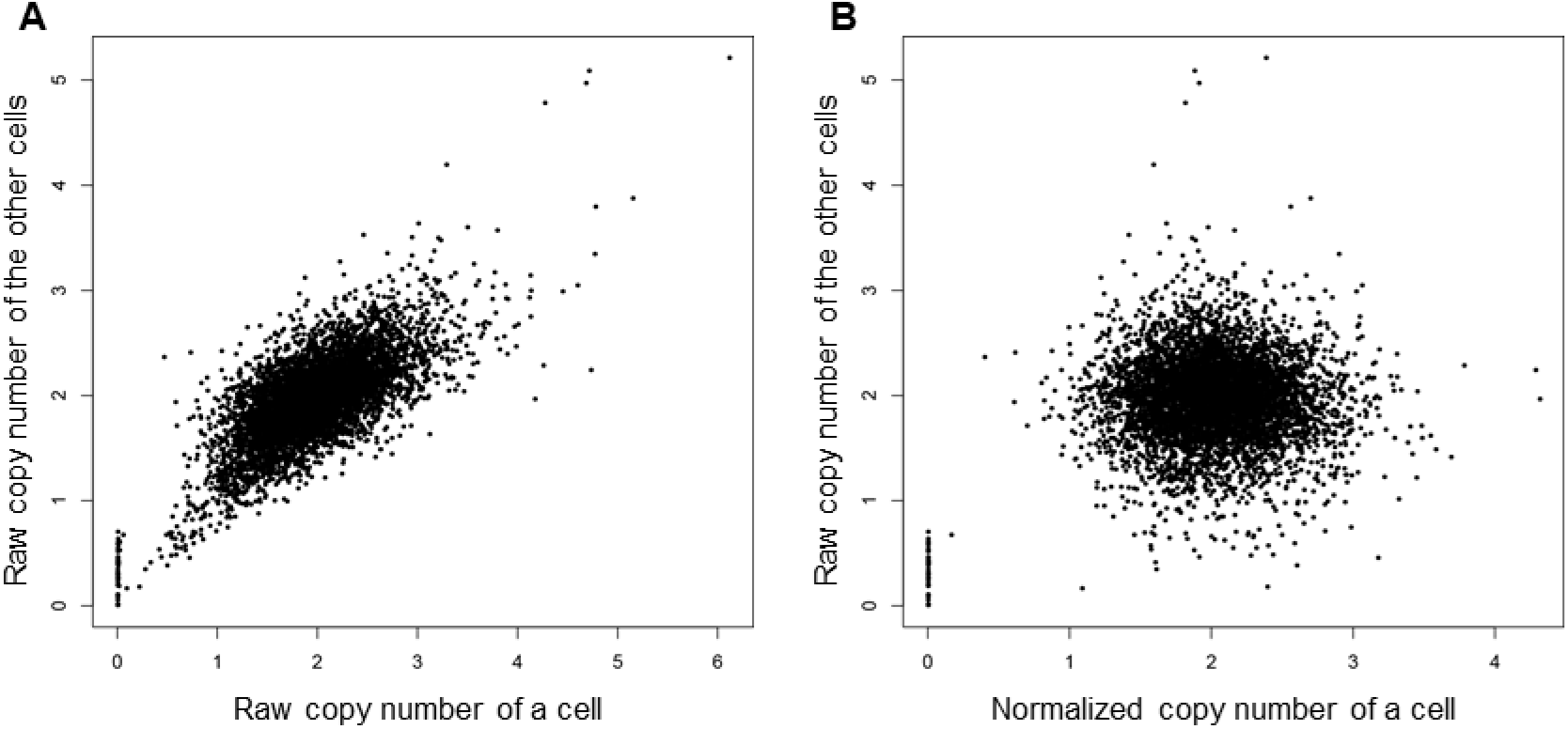
Correlation in copy number estimation between a cell and the other cells of the same batch before (A) and after (B) normalizing locus-specific amplification bias. This example was generated using a cell amplified with MDA (SRA id: SRR2141574).

**Figure 5.**
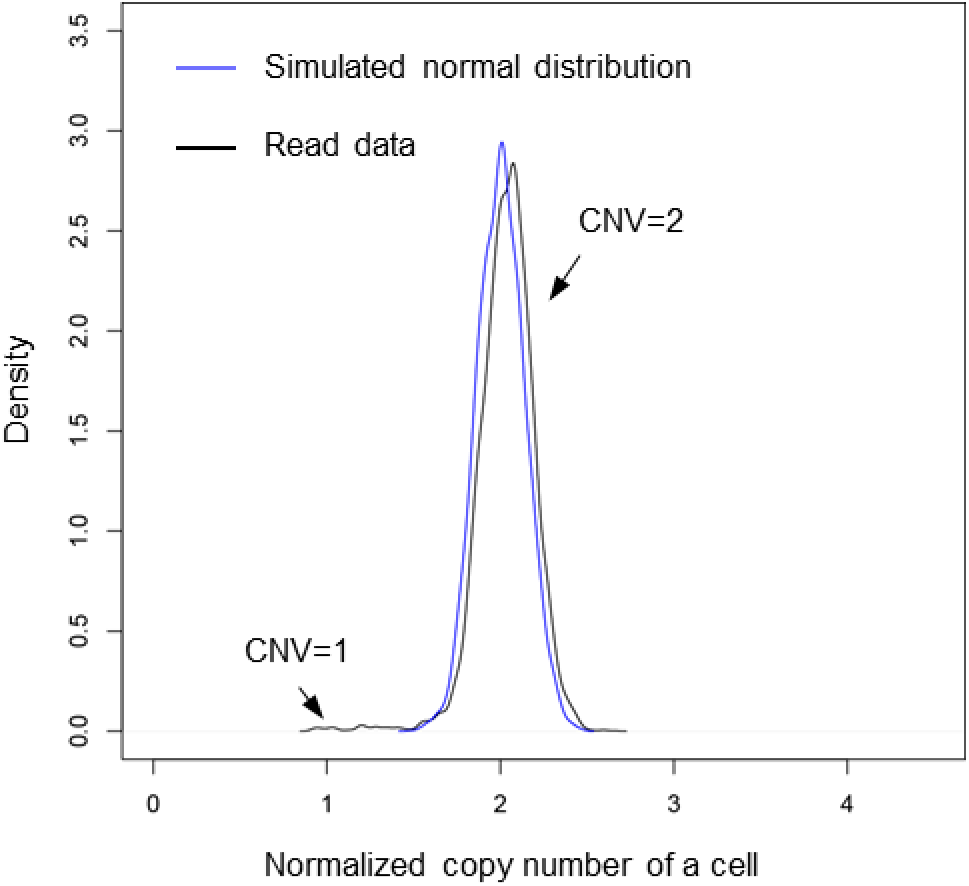
Inferring copy number variation by comparing the observed with a simulated distribution. This example was generated using a cell amplified with MDA (SRA id: SRR2141574).

SCCNV was applied to three publicly available SCWGS datasets [2, 5, 8]. The three datasets included SCWGS of 60 single human fibroblasts or neurons amplified using 8 different protocols, i.e., DOP-PCR (Sigma), Rubicon, MALBAC (Yikon), LIANTI, and MDA (including Qiagen, GE, Lodato et al’s MDA and SCMDA). Sequence alignment was performed using BWA and GATK (Supplementary Material) [9, 10]. Raw sequencing data of each sample (cell and bulk DNA) were obtained from SRA database and subjected to quality control using FastQC [11] and trimming using Trim Galore [12] with default parameters. Then they were aligned to human reference genome (version hg19) using BWA MEM [9]. PCR duplications were removed using picard tools [13]. The alignments were subjected to indel realignment and basepair recalibration using GATK (version 3.5) [10]. Reads with mapQ<30 were discarded. Numbers of reads per bin of each sample was calculated using samtools [14].

As shown in Table 1, single cells amplified using DOP-PCR, Rubicon, MALBAC and LIANTI are generally of sufficiently good quality to determine CNVs: CNVs can be inferred for up to 100% of the genome. Among these methods, LIANTI performed the best. This is consistent with a previous report [5]. MDA-based methods were found to perform much less as compared to the others (up to 68.8% of the genome), although they suffer much less from artifactual SNVs [2].

**Table 1.**
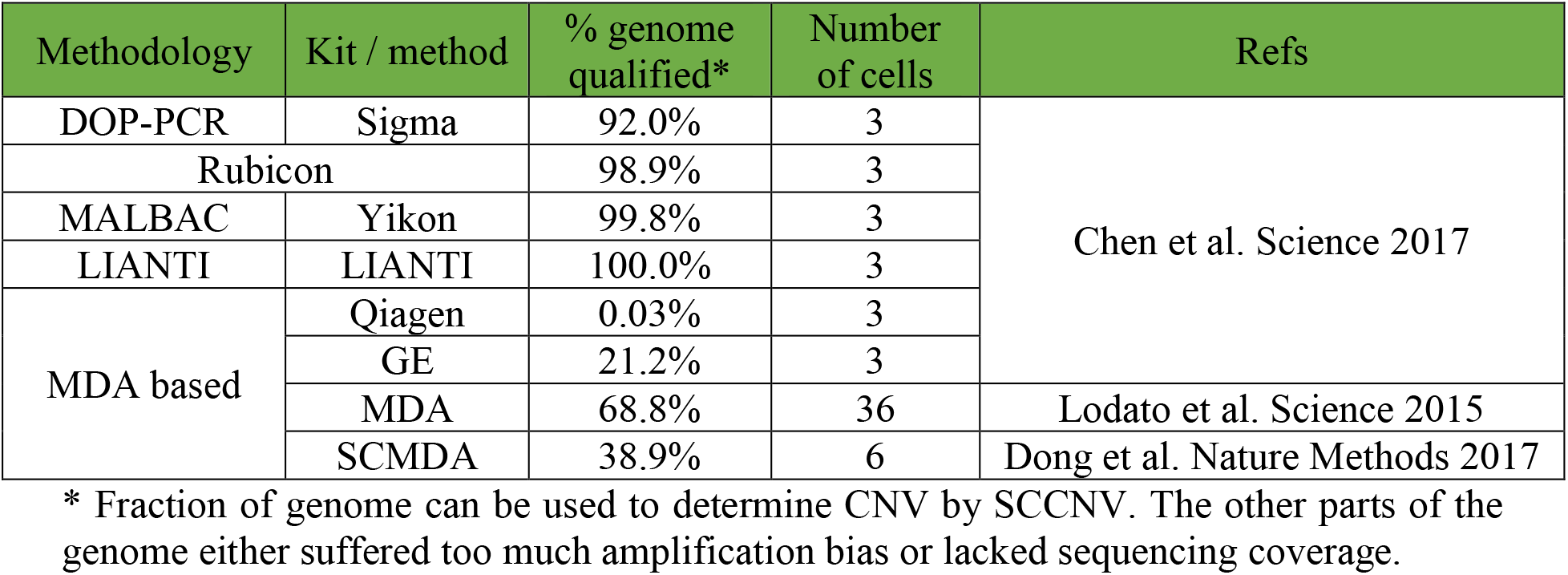
A summary of data used for CNV analyses.

## Conclusions

We developed SCCNV to identify copy number variations from whole-genome amplified single cells. We demonstrated its performance using some of the recent SCWGS datasets generated with 8 different amplification protocols.

## Availability and requirements

Project name: SCCNV.

Project home page: https://github.com/biosinodx/SCCNV.

Operating system(s): Linux or MAC OS.

Programming language: Python.

Other requirements: Python 2 or 3, Python module numpy, bedtools,

License: GNU GPL version 3 or later.

Any restrictions to use by non-academics: No.

## List of abbreviations

SCWGS: single-cell whole-genome sequencing
CNV: copy number variation
WGA: whole-genome amplification

## Declarations

### Availability of data and material

SCCNV is freely available at https://github.com/biosinodx/SCCNV.

### Authors’ contributions

XD, LZ and JV conceived the study. XD and TW developed the method. XD and XH analyzed the data. XD, LZ and JV wrote the manuscript.

## Acknowledgements

Not applicable.

